# Leukocyte counts based on site-specific DNA methylation analysis

**DOI:** 10.1101/150110

**Authors:** Joana Frobel, Tanja Božić, Michael Lenz, Peter Uciechowski, Yang Han, Reinhild Herwartz, Klaus Strathmann, Susanne Isfort, Jens Panse, André Esser, Carina Birkhofer, Uwe Gerstenmaier, Thomas Kraus, Lothar Rink, Steffen Koschmieder, Wolfgang Wagner

## Abstract

The composition of white blood cells is usually assessed by histomorphological parameters or flow cytometric measurements. Alternatively, leukocyte differential counts (LDCs) can be estimated by deconvolution algorithms for genome-wide DNA methylation (DNAm) profiles. We identified cell-type specific CG dinucleotides (CpGs) that facilitate relative quantification of leukocyte subsets. Site-specific analysis of DNAm levels by pyrosequencing provides similar precision of LDCs as conventional methods, whereas it is also applicable to frozen samples and requires only very small volumes of blood. Furthermore, we describe a new approach for absolute quantification of cell numbers based on a non-methylated reference DNA. Our “Epi-Blood-Count” facilitates robust and cost effective analysis of blood counts for clinical application.

## Introduction

Leukocyte differential counts (LDCs) give valuable diagnostic insights for various systemic and malignant diseases. It is among the most frequently requested laboratory tests in hematological diagnostics (Buttarello & Plebani, 2008). LDCs can be determined by microscopic evaluation and manual counting. However, since the advent of automated cell counters, LDCs are particularly analyzed by flow cytometric technologies (Estridge & Reynolds, 2011; Koepke et al, 2007; Roussel et al, 2010). Such automated analyzers sense electrical impedance, optical light scattering properties, or fluorescence signal intensities (Briggs et al, 2014; Cherian et al, 2010; Roussel et al, 2012). Fluorescent staining of specific epitopes is the gold standard for definition of lymphocyte subsets. However, immunophenotypic analysis is costly, relatively labor-intensive and not trivial to standardize. Furthermore, all of the aforementioned methods are only applicable to fresh blood samples – it is not possible to freeze samples for shipment or later analysis, e.g. when blood samples cannot be immediately processed (Briggs et al, 2014; Buoro et al, 2016; Zini, 2014). Recently, gene expression profiles (Abbas et al, 2009; Gong et al, 2011; Newman et al, 2015; Shen-Orr & Gaujoux, 2013) as well as epigenetic profiles (Accomando et al, 2014; Houseman et al, 2012) have been used to deconvolute the cellular composition in whole blood. Such alternative approaches might overcome some of the limitations of the well-established state of the art procedures for LDCs.

DNA-methylation (DNAm) represents the best understood epigenetic modification, and it is directly linked to developmental processes (Smith & Meissner, 2013). Methyl groups can be added to the 5^th^ carbon atom of cytosines, predominantly in a cytosine-guanine dinucleotide (CpG) context (Lister et al, 2009). Many studies demonstrated that genome-wide DNAm profiles can be used to estimate LDCs (Adalsteinsson et al, 2012; Houseman et al, 2015; Jaffe & Irizarry, 2014; McGregor et al, 2016; Waite et al, 2016). In fact, DNAm patterns have many advantages compared to immunophenotypic analyses: i) DNAm is directly linked to cellular differentiation; ii) it facilitates absolute quantification at single base resolution (ranging from 0 to 100% DNAm); iii) every cell has only two copies of DNA and hence the results can be easily extrapolated to the cellular composition (in contrast to RNA, which can be highly overexpressed in small subsets); and iv) DNA is relatively stable: it can be isolated from lysed or frozen cells and shipped at room temperature for further analysis. Deconvolution algorithms to estimate the cellular composition in heterogeneous tissues are usually based on a-priori reference datasets of cell type-specific DNAm patterns (Teschendorff et al, 2017). In principle, it is also possible to train reference-free algorithms, but since the composition of blood is relatively well known, reference-based algorithms appear to be advantageous (Zheng et al, 2017). So far, the available algorithms for LDCs are based on many CpGs of the 450k Illumina BeadChip microarray – and these methods would need to be reestablished for the newer EPIC array, whole genome bisulfite sequencing (WGBS), or reduced representation bisulfite sequencing (RRBS) data (BLUEPRINT consortium, 2016). Either way, all of these profiling procedures are hardly applicable in daily clinical routine.

In this study, we followed the hypothesis that even site-specific analysis of DNAm at individual CpG sites can be utilized to determine LDCs. The results of our “Epi-Blood-Count” reveal similar precision as conventional methods. Furthermore, we conceived a method to quantify absolute cell counts based on DNAm patterns.

## Results

### Identification of individual CpG sites to discern hematopoietic subsets

For selection of candidate CpGs we utilized DNAm data of purified granulocytes, CD4+ T cells, CD8+ T cells, B cells, NK cells, and monocytes (GSE35069) that has been published by Reinius and coworkers (Reinius et al, 2012). For each of these cell types we selected CpG sites that facilitate best discrimination based on the following two criteria (Fig. 1 A): i) highest difference in mean β-value of the subset and the mean β-value of all other hematopoietic cell types; and ii) low variance of β-values across different samples of the corresponding subset. This analysis was initially performed for granulocytes (Fig. 1 B and C) and then repeated for the other cell types (Figs. S1 and S2). Furthermore, we combined DNAm profiles of T cells, B cells, and NK cells to identify CpGs that reflect the entire lymphocyte population; and of CD4+ and CD8+ T cells as a surrogate for the entire T cell population. Best performing CpG sites were validated on a second dataset of purified leukocyte subsets (E-MTAB-2145; Fig. 1 D) (Zilbauer et al, 2013).

**Figure 1.**
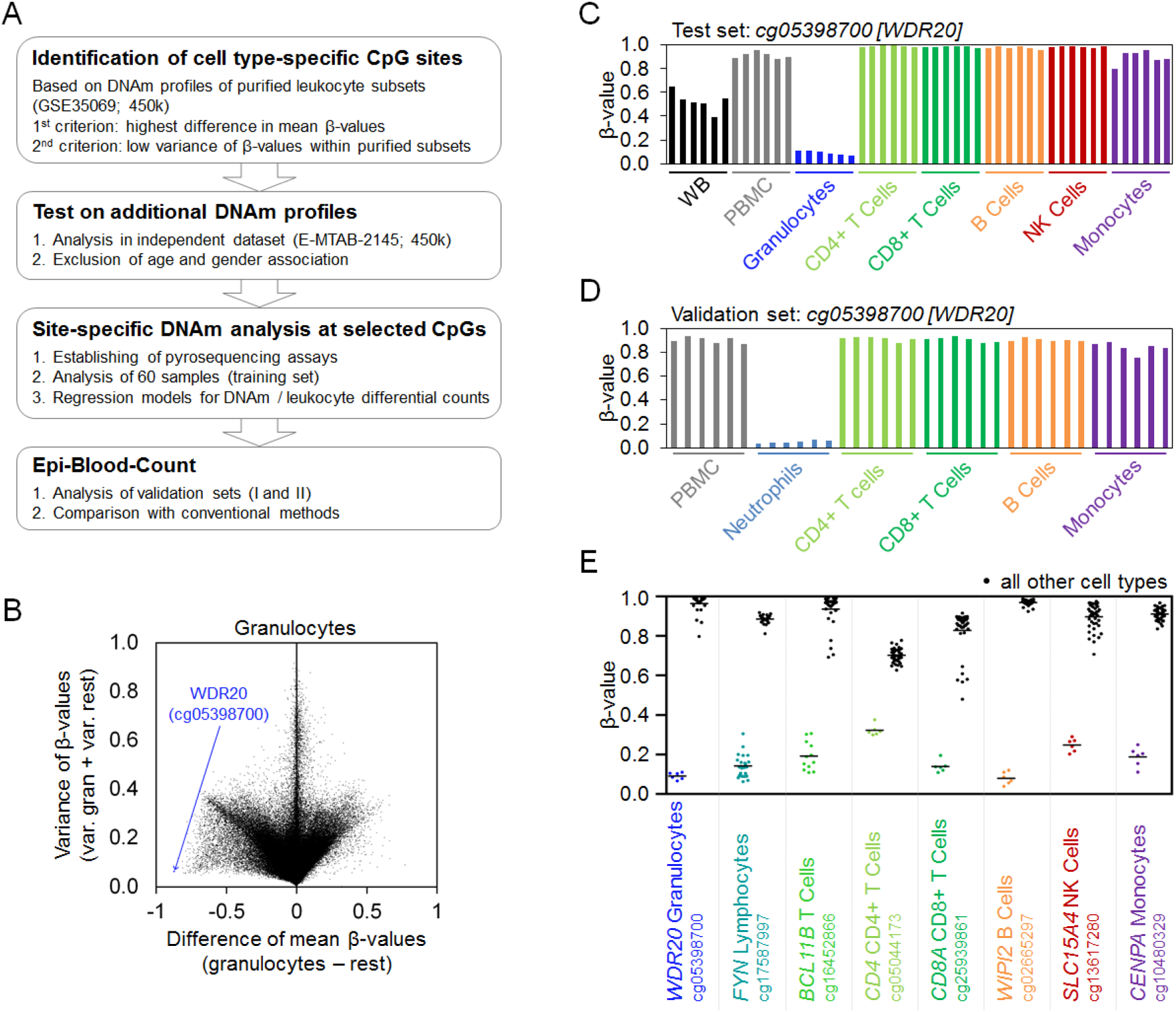
Selection of cell type-specific CpG sites for the Epi-Blood-Count. (A) Schematic presentation of the work-flow that led to selection of candidate CpG sites for individual hematopoietic subsets. (B) Scatterplot to exemplarily depict selection criteria for the CpG site for granulocytes (cg05398700): i) high difference between mean β-values in granulocytes (gran.) and the mean β-values of all other cell types (rest) in DNAm profiles of purified cell types (GSE35069) (Reinius et al, 2012); and ii) a low variance (var.) of β-values within the granulocytes and within the other hematopoietic subsets. (C) Differences in β-values across blood cell types are exemplarily demonstrated for cg05398700 in reference dataset of Reinius et al. (GSE35069) (Reinius et al, 2012), and (D) in the validation set of Zilbauer et al. (E-MTAB-2145) (Zilbauer et al, 2013). (E) In analogy, the other cell type-specific CpGs of the Epi-Blood-Count were selected. DNAm levels are depicted for each cell type-specific CpG in comparison to all other hematopoietic subsets (GSE35069).

For granulocytes we selected a CpG site in the gene WD repeat domain 20 (*WDR20*; cg05398700). Notably, CpGs with highest discriminatory power for CD4+ T cells and CD8+ T cells are linked to the genes *CD4* (cg05044173) and *CD8A* (cg25939861), respectively. Furthermore, the selected CpG site for lymphocytes was in the FYN Proto-Oncogene (*FYN;* cg17587997); for T cells in B-cell lymphoma/leukemia 11B (*BCL11B;* cg16452866) associated with B cell malignancies; for B cells in WD repeat domain, phosphoinositide interacting 2 (*WIPI2;* cg02665297) that is involved in maturation of phagosomes; for NK cells in solute carrier family 15 member 4 (*SLC15A4;* cg13617280) that has been implicated in systemic lupus erythematosus; and for monocytes in centromere protein A (*CENPA*; cg10480329; Fig. 1 E). Thus, our straight-forward procedure identified CpGs that are associated with genes of relevant function in the corresponding cell types.

Subsequently, we analyzed if site-specific DNAm measurements at our candidate CpGs correlate with the fractions of hematopoietic subsets in blood. To this end, we established pyrosequencing assays for the selected CpG sites and analyzed 60 peripheral blood samples. The percentage of leukocytes – as determined with a Sysmex XN-9000 hematology analyzer – correlated very well with DNAm at the respective CpG sites for granulocytes (R = −0.91), lymphocytes (R = −0.91), and monocytes (R = −0.74). Furthermore, immunophenotypic lymphocyte subset enumerations correlated for T cells (R = −0.73), CD4+ T cells (R = −0.41), CD8+ T cells (R = −0.88), B cells (R = −0.66), and to a lesser extent for NK cells (R = −0.30; Fig. S3). The correlation coefficients are negative because all selected CpGs were hypomethylated in the corresponding subsets. Thus, DNAm at unique candidate CpGs reflects the frequency of corresponding cell types in blood. Furthermore, we excluded that our candidate CpGs were associated with aging or gender (Fig. S4). There was only a moderate age-associated decline in DNAm at the CpGs for CD4+ T cells and monocytes, and this may correspond to the commonly observed changes upon aging (Gruver et al, 2007; Melzer et al, 2015).

### Deconvolution of granulocytes, monocytes, and lymphocytes with three CpGs

The fractions of granulocytes, monocytes, and lymphocytes are routinely determined with automated analyzers or manual microscopic counting. To evaluate if such results can be recapitulated by the three corresponding CpGs we measured DNAm levels in 44 independent blood samples by pyrosequencing. Initially, the percentages of cells were simply calculated based on the linear regression formulas of the subsets in the training set (Fig. S3). In comparison to measurements of the Sysmex XN-9000 analyzer, the linear regression models based on three CpGs revealed extremely high correlation (R = 0.99 across all cell types). The mean absolute deviation (MAD) was only 3.2% for granulocytes, 2.2% for lymphocytes, and 1.4% for monocytes (Fig. S5).

Alternatively, we integrated the DNAm levels of the three CpGs into a non-negative least-squares (NNLS) linear regression model. This model was trained on 60 blood samples of the training set and is subsequently termed “Epi-Blood-Count”. The NNLS linear regression approach does not depend on an a-priori database of cell type-specific DNAm reference profiles for the selected CpG sites. Either way, the estimated DNAm levels based on deconvolution of 60 blood samples of the training set were very similar to the β-values of DNAm profiles of purified subsets (Reinius et al, 2012) (Fig. S6). This approach gave similar accuracies as the linear models based on individual CpGs for granulocytes, lymphocytes, and monocytes (Fig. 2 A and B).

**Figure 2.**
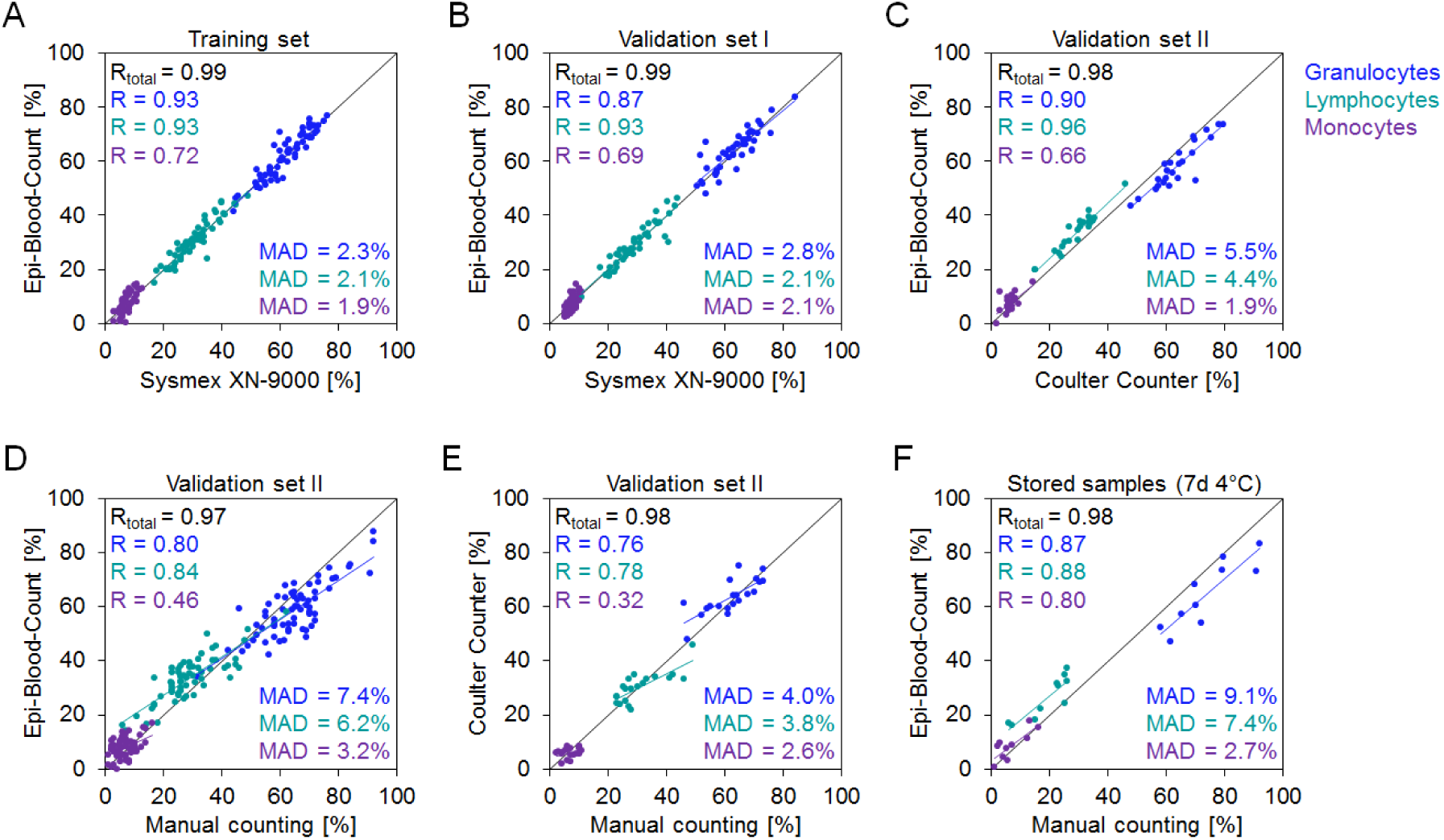
Epi-Blood-Count of granulocytes, lymphocytes, and monocytes. Peripheral blood samples of a training set (A) (n = 60) and of an independent validation set (B) (n = 44) were analyzed with a Sysmex XN-9000 hematology analyzer and DNAm was analyzed by pyrosequencing at the three CpGs for granulocytes, lymphocytes, and monocytes. Furthermore, performance of the Epi-Blood-Count was compared with (C) a Coulter Counter (ACT II Diff Counter from Beckmann Coulter; n = 24) and/or (D) microscopic analysis of blood smears and manual counting by a trained operator (n = 66). (E) The correlation of Coulter Counter results and manual counting was compared (n = 20). (F) To determine how the results of the Epi-Blood-Count are affected by long-term storage (without freezing), we used an additional set of blood samples (n = 10) that was initially analyzed by manual counting (at day 0) and by DNAm analysis at the three relevant CpGs after storage for seven days at 4°C. Pearson correlation coefficient (R) and mean absolute deviation (MAD) between the measurements were calculated for each cell type.

Our Epi-Blood-Count was trained on cell counts that were determined with the Sysmex XN-9000 analyzer – however, there are notoriously differences between cell counting systems (Buttarello & Plebani, 2008; Estridge & Reynolds, 2011). Therefore, we applied the Epi-Blood-Count on a second validation set (in total 70 blood samples) that were either measured with a Coulter Counter (ACT II Diff Counter, Beckman Coulter; n = 24), and/or by manual counting of blood smears by highly specialized laboratory staff (n = 66). Coulter Counter results revealed very high correlation with the Epi-Blood-Count, albeit granulocytes and lymphocytes were – on average – underestimated by 4.4% and overestimated by 5.5%, respectively, indicating that there might be a systemic deviation between the two analyzers (Fig. 2 C). The correlation between manual blood counts and Epi-Blood-Count was slightly lower (Fig. 2 D), but direct comparison of Coulter Counter results and manual counting revealed lower correlations, too (Fig. 2 E). Furthermore, we have exemplarily analyzed if site-specific DNAm analysis for Epi-Blood-Count is also feasible with other techniques: MassARRAY analysis is based on mass spectrometric analysis of DNA fragments and might be advantageous for high throughput analysis in a 384-well format. Measurements of DNAm levels by pyrosequencing and MassARRAY analysis correlated and Epi-Blood-Count based on MassARRAY technology is in principle feasible, but there was a systematic offset that needs to be taken into consideration (Fig. S7).

Long-term storage of blood affects LDCs (Gulati et al, 2002). We therefore tested if the Epi-Blood-Count is also applicable after storage of blood samples for seven days at 4°C. Overall, the results correlated with manual counting at day 0, but the average numbers of granulocytes and lymphocytes were underestimated by 9.1% or overestimated by 7.2%, respectively (Fig. 2 F). With automated analyzers a similar shift was already observed upon storage of blood samples for 72 hours (Joshi et al, 2015). Furthermore, we anticipated that it is possible to freeze fresh blood samples and store them for months without affecting the results of the Epi-Blood-Count. To test this, we used a third validation set (n = 41) that was either analyzed by Epi-Blood-Count in aliquots of fresh blood or after being frozen for three months. DNAm levels at the relevant CpGs were not affected by long-term cryopreservation (Fig. S8). Another advantage of the Epi-Blood-Count is that it is applicable to very low amounts of DNA. We have exemplarily used this approach to measure cell-free circulating DNA (cfDNA) in serum. The results indicated that cfDNA in serum is particularly derived from granulocytes, which indeed have a very short half-life (Summers et al, 2010) (Fig. S9).

### Additional classification of lymphocyte subsets

Subsequently, we analyzed whether the Epi-Blood-Count can be further extended to classify subsets of lymphocytes. To this end, the DNAm levels at the candidate CpGs for B cells, NK cells, CD4+ T cells, CD8+ T cells, and T cells (combination of CD4+ and CD8+ T cells was used as surrogate for the entire T cell population) were analyzed by pyrosequencing in the 60 blood samples of the training set. Since the percentages of these subsets correlated well with DNAm levels (Fig. S3) it was again possible to determine the LDCs based on linear regression formulas (Fig. S10). Alternatively, the corresponding immunophenotypic measurements of the training set were imputed to generate NNLS regression models for either five CpGs (CD4+ and CD8+ T cells combined) or six CpGs (separate analysis of CD4+ and CD8+ T cells). With this deconvolution approach the estimated DNAm levels for each hematopoietic subset closely resembled the β-values of the purified subsets in DNAm profiles (Reinius et al, 2012) (Fig. S11 A and B). The five CpG and six CpG Epi-Blood-Count models were then tested on the training set (Fig. S11 C and D) and on two independent validation sets (Fig. 3 and Fig. S11 E and F). Immunophenotypic analysis and Epi-Blood-Count revealed a clear correlation: across all cell types the correlation coefficients for the five CpG and six CpG models were R = 0.99 and R = 0.98 with average MADs of 2.8% and 3.1%, respectively. Furthermore, the measurements were even relatively stable after storage of blood samples at 4°C for seven days without fixation (Fig. S12).

**Figure 3.**
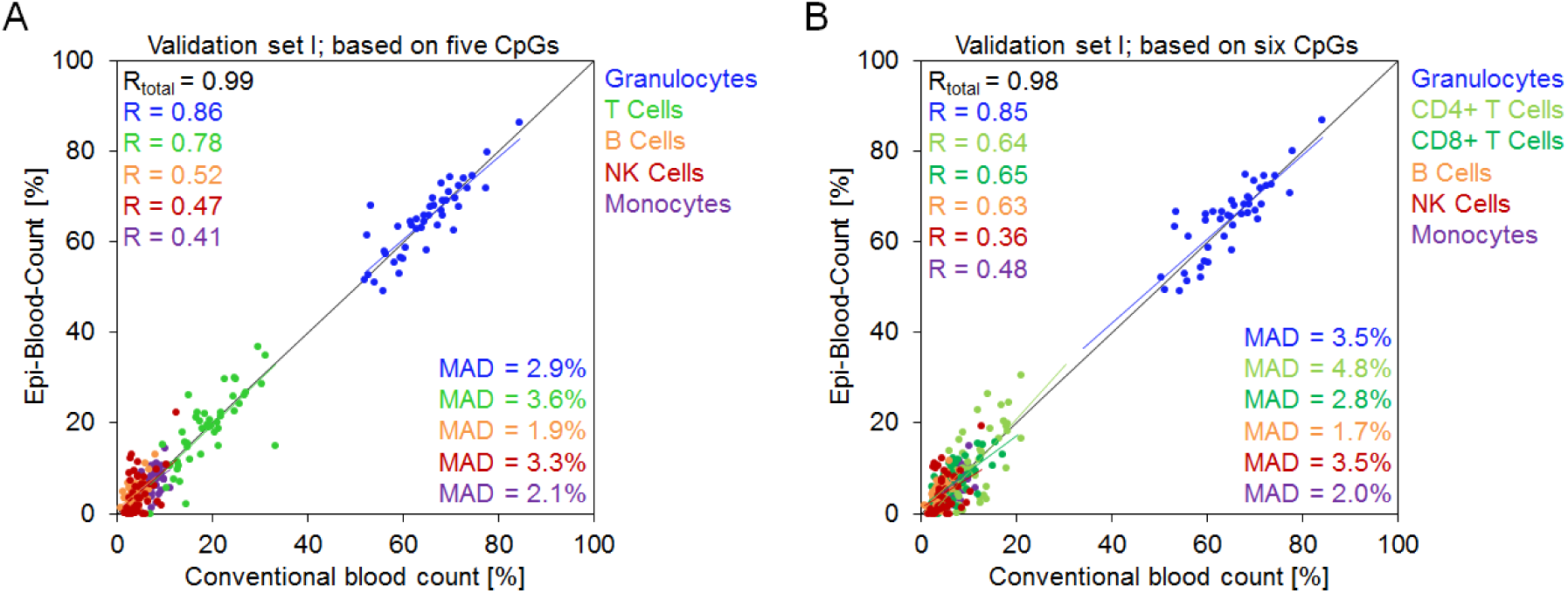
Leukocyte differential counts with five and six CpG sites. Leukocyte differential counts were determined in the validation set I by combination of Sysmex XN-9000 hematology analyzer and immunophenotypic analysis with FACSCalibur (n = 44). The conventional LDCs were compared with Epi-Blood-Count results that were either determined based on DNAm analysis by pyrosequencing of (A) five CpGs (including a CpG site for all T cells), or (B) six CpGs to further discriminate between CD4+ and CD8+ T cells. The models for the Epi-Blood-Count (NNLS) were previously generated on an independent training set. R = Pearson correlation coefficient; MAD = mean absolute deviation.

### Epigenetic analysis of cell numbers

Conventional flow cytometric methods facilitate measurement of absolute cell numbers per μl blood, whereas deconvolution of gene expression profiles and DNAm profiles provides only proportionate cell fractions. We reasoned that quantification of cell numbers based on DNAm levels would be feasible if samples were spiked with a suitable reference DNA of known concentration (Fig. 4 A). To this end, we identified three CpG sites that are consistently highly methylated (β-value > 0.975) across all DNAm profiles of hematopoietic subsets and in DNAm profiles of whole blood of healthy individuals, leukemia or lymphoma patients (Fig. S13). The selected CpG sites were within LSM family member 14B (*LSM14B*; cg06096175), zinc finger CCCH-type containing 3 (*ZC3H3*, cg25834632) and a CpG site not associated with any gene (cg09414987). The corresponding sequences were cloned into plasmids and amplified in *E. coli* to obtain non-methylated reference DNAs.

**Figure 4.**
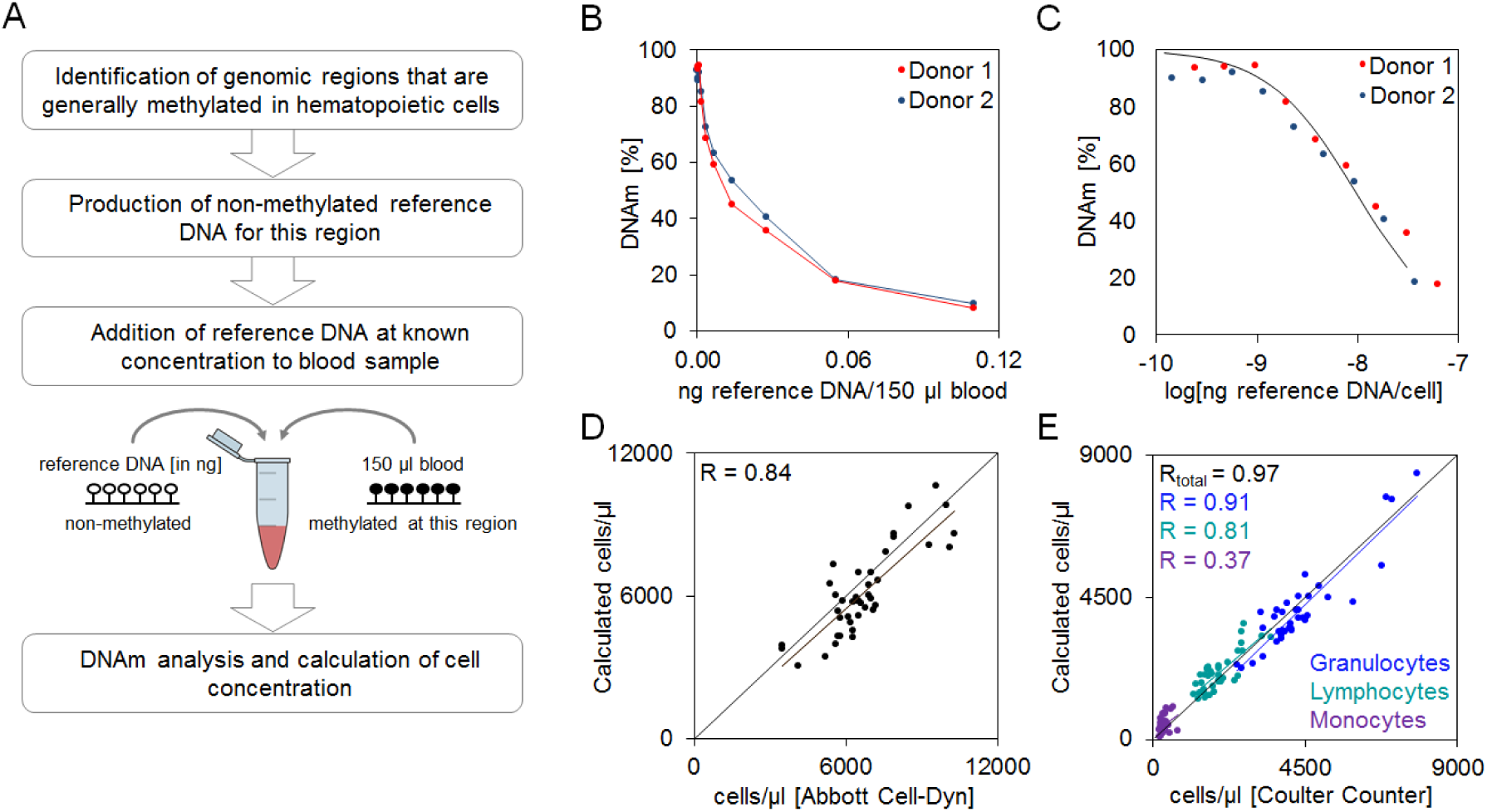
Quantification of cells based on DNA methylation. (A) Schematic presentation of cellular quantification based on DNAm levels with a non-methylated reference DNA. (B) Two blood samples (donor 1 and 2) were mixed with a serial dilution of the reference plasmid comprising the non-methylated sequence of *LSM14B* (0.0002 ng to 0.1100 ng). DNAm levels (analyzed by pyrosequencing) continuously declined with higher concentrations of reference DNA. (C) If the results were plotted as logarithmic ratio of reference DNA [ng] per cell (determined with the Abbott Cell-Dyn), there was an almost linear association in the DNAm range between 20% and 80%. Notably, the observed DNAm levels closely resembled the expected calculated DNAm levels (black curve). (D) Calculated cell numbers based on the reference plasmid *LSM14B* clearly correlated with cell numbers determined with the Abbott Cell-Dyn analyzer (n = 41; validation set III). (E) Furthermore, epigenetic quantification could be combined with epigenetic LDCs: Cell numbers for granulocytes, lymphocytes, and monocytes correlated with cell numbers determined with a Coulter Counter (n = 38; validation set IV; MAD = mean absolute deviation).

Initially, we analyzed serial dilutions of reference DNA (*LSM14B*) in two independent peripheral blood samples and the DNAm levels declined with higher proportions of reference DNA (Fig. 4 B). The predicted DNAm levels based on mathematical calculation with all known parameters correlated very well with the measured DNAm levels, indicating that the method is robust for cellular quantification (Fig. 4 C). The precision of this approach is particularly high if DNAm levels range between 20% and 80% – if copy numbers of reference DNA and genomic DNA are similar. To increase this range we used the other two reference DNAs at higher and lower concentration, respectively (Fig. S14).

Subsequently, we mixed 150 μl of frozen blood samples (n = 41; validation set III) with our *LSM14B* reference DNA and analyzed DNAm levels at the relevant CpG site by pyrosequencing. The calculated cell numbers correlated well with cell counts that were automatically measured in fresh blood (R = 0.84; Fig. 4 D). Furthermore, combined epigenetic analysis of relative LDCs with absolute cell numbers correlated with measurements of an automated hematology analyzer for individual hematopoietic subsets (n = 38; validation set IV; R = 0.97; Fig. 4 E). This method is in principle also applicable for MassARRAY technology (Fig. S15).

## Discussion

Analysis of DNAm patterns in blood holds enormous diagnostic potential, which has so far hardly been utilized. We demonstrate that site-specific analysis at individual CpG sites facilitates relative quantification of leukocyte subpopulations. Furthermore, we describe a new method of using a non-methylated reference DNA to determine absolute cell numbers based on DNAm.

Analytic performance is traditionally evaluated by precision (or random error), accuracy (or systematic error), and clinical sensitivity. It was somewhat unexpected that analysis of DNAm at only few individual CpG sites reached similar precision and accuracy as the well-established conventional methods (Buttarello & Plebani, 2008; Estridge & Reynolds, 2011). However, it remains to be demonstrated if this is also applicable to clinical samples. Diseases – particularly hematopoietic malignancies – have major impact on the epigenetic makeup and hence it will be important to determine how the epigenetic blood signatures are affected by specific diseases. In this regard, it may be advantageous to consider additional genomic regions for hematopoietic subsets to validate that cell type-specific CpGs are coherently modified. However, analysis of a higher number of CpGs is always a trade-off with regard to costs and time.

The previously published algorithms for LDCs are based on DNAm profiles that were generated with Illumina BeadChip microarrays (Accomando et al, 2014; Houseman et al, 2012; Teschendorff et al, 2017). Such larger signatures have the big advantage to combine a multitude of CpGs, which generally increases the precision of epigenetic signatures (Koestler et al, 2013). On the other hand, the precision of DNAm levels at individual CpGs is higher in pyrosequencing data as compared to β-values on Illumina BeadChips (BLUEPRINT consortium, 2016; Lin et al, 2016). Furthermore, analysis of genome-wide DNAm profiles takes longer and it is relatively expensive. This is also the reason why the number of available DNAm profiles with matched flow cytometric analysis is still relatively low. Reinius et al. provided flow cytometric analysis for six DNAm profiles (Reinius et al, 2012) and Absher and colleagues provided 44 DNAm profiles with conventional LDCs (Absher et al, 2013). Notably, the precision of genome-wide algorithms on these datasets was similar to the performance of the Epi-Blood-Count in our cohorts (Tables S1 and S2) (Waite et al, 2016). Furthermore, Koestler and coworkers compared their predictions of cell types with complete blood counts that were analyzed on an Advia 70 hematology system and the correlation for monocytes (R = 0.60) and lymphocytes (R = 0.61) was not better than our 3 CpG Epi-Blood-Count (Koestler et al, 2013). A recent study indicated that non-constrained methods, such as support vector regressions or robust partial regression, might further improve the accuracy of deconvolution methods for DNAm profiles (Teschendorff et al, 2017). Either way, site-specific analysis of individual cell type-specific CpGs is better applicable to daily routine in clinical diagnostics.

Our proof-of-concept study should be further developed to address additional cell types in the future. So far, the Epi-Blood-Count does not consider eosinophils, basophils, immature granulocytic precursors, or more specialized lymphocyte subsets such as naïve, memory, or regulatory T cells. It has been demonstrated that the percentage of these additional cell types can be estimated based on gene expression or DNAm data (Newman et al, 2015; Waite et al, 2016). Furthermore, we expect that it is possible to integrate CpGs that are indicative for blasts, atypical lymphocytes, and hematopoietic progenitors for extended differential counts. For rare cell types, it will be more challenging to identify candidate CpGs that reflect the percentage in peripheral blood and corresponding DNAm profiles of purified subsets need to be available. It is conceivable that alternative methods for DNAm analysis, such as barcoded bisulfite amplicon sequencing (BBA-seq) (Franzen et al, 2016; Masser et al, 2013) or digital PCR (Weisenberger et al, 2008), may ultimately pave the way for more sensitive deconvolution of rare subsets. Furthermore, analysis of neighboring CpG sites of the same amplicon may increase robustness as described for detection of circulating tumor DNA (Lehmann-Werman et al, 2016).

Relative quantification of cell types is of relevance for clinical diagnostics, but it is very important to determine absolute cell numbers as well (Buttarello & Plebani, 2008; Koepke et al, 2007). In this study, we describe an entirely new approach for cellular quantification based on DNAm levels, which is based on addition of a non-methylated reference sequence of known concentration. In analogy, quantification of cell numbers has been established in flow cytometry by addition of beads as quantitation standards (Montes et al, 2006). Comparison with other established methods for cell counting indicated that the precision of our DNAm based approach is similar (Cadena-Herrera et al, 2015). Addition of non-methylated reference DNA would even be applicable before analysis of genome-wide DNAm profiles by Illumina BeadChips, WGBS, and RRBS – and thereby the quantification approach might actually be combined with larger deconvolution algorithms based on multiple CpGs.

Site-specific analysis of DNAm levels at individual CpGs can reflect the relative abundance of hematopoietic subsets. Immunophenotypic analysis is based on individual cell type-specific epitopes, too. Notably, several candidate CpGs of the Epi-Blood-Count are related to the same genes addressed in immunophenotypic analysis. Our Epi-Blood-Count has various advantages over the well-established conventional methods: i) blood can be frozen after sampling for long-term storage, shipment, and subsequent analysis; ii) it is applicable to relatively small volumes of blood (less than 100 μl, whereas at least 700 μl are required for immunophenotypic analysis); and iii) DNAm levels at individual CpGs provide an absolute measure that may facilitate better standardization between labs than immunophenotypic analysis by flow cytometry. In combination with a non-methylated standard sequence it is possible determine absolute cell numbers based on DNAm. It is, however, unlikely that epigenetic analysis of LDCs will completely replace the conventional cell counters, because it cannot address erythrocytes and thrombocytes, which hardly comprise DNA.

## Materials and Methods

### Selection of candidate CpGs

For selection of cell type-specific CpG sites we used DNAm profiles of purified leukocyte subsets that were generated on the Illumina Infinium HumanMethylation450 BeadChip platform (Gene Expression Omnibus ID: GSE35069) (Reinius et al, 2012). We utilized β-values, ranging from 0 to 1, which provide a measure for each CpG site represented on the array that roughly corresponds to the percentage of DNAm. CpG sites on X and Y chromosomes were excluded. Candidate CpGs were selected based on i) highest difference between the mean β-value of one purified leukocyte subset and the mean β-value of all other subsets; and ii) low variation of β-values within purified subsets. Best performing CpG sites were further evaluated on an independent dataset of purified leukocyte subsets provided on the Array Express database (E-MTAB-2145) (Zilbauer et al, 2013).

For selection of CpGs for cellular quantification we identified genomic regions that were consistently methylated across all hematopoietic subsets. To this end, we used the following DNAm profiles that were all analyzed on the HumanMethylation450 BeadChip: i) purified leukocyte subsets: GSE35069 (Reinius et al, 2012), and E-MTAB-2145 (Zilbauer et al, 2013); ii) whole blood from healthy donors: GSE32148 (Harris et al, 2012), GSE41169 (Horvath & Levine, 2015); and iii) DNAm profiles of blood disorders such as acute myeloid leukemia: TCGA (The Cancer Genome Atlas Research Network, 2013), GSE58477 (Qu et al, 2014), GSE62298 (Ferreira et al, 2016); myelodysplastic syndrome: GSE51758 (Zhao et al, 2014); B cell lymphoma: GSE37362 (Asmar et al, 2013); acute lymphoblastic leukemia: GSE69954 (Borssen et al, 2016)). Mean β-values were calculated for each dataset and all CpGs. Subsequently, we selected three candidate CpG sites (cg06096175 (*LSM14B);* cg25834632 (*ZC3H3*); cg09414987 (no associated gene)) that were consistently highly methylated in each dataset (β-value > 0.975).

### Blood samples

Peripheral blood samples for the training set (n = 60) and for validation set I (n = 44) were obtained from the HELPcB program (Health Effects in High-Level Exposure to PCB) (Schettgen et al, 2012). The study was approved by the local ethics committee of the RWTH Aachen University (EK 176/11). Peripheral blood samples for validation set II (n = 70), validation set III (n = 41), and validation set IV (n = 38), as well as serum samples (n = 18) were obtained from the Department of Hematology, Oncology, Hemostaseology, and Stem Cell Transplantation and from the Department of Transfusion Medicine according to the guidelines specifically approved by the local ethics committee of the RWTH Aachen University (EK 099/14).

### Conventional analysis of blood counts

Blood samples from the HELPcB program were analyzed with the Sysmex XN-9000 hematology analyzer (Sysmex Deutschland GmbH, Norderstedt, Germany). Immunophenotyping was performed as described before (Haase et al, 2016). In brief, EDTA anti-coagulated whole blood was incubated for 20 minutes at room temperature with fluorescently labeled antibody pairs (CD3/CD4, CD3/CD8, CD3/CD19, CD3/CD16+CD56) and isotype matched controls (IgG1 FITC/IgG2a PE, all from Becton Dickinson, Heidelberg, Germany). Erythrocytes were lysed with BD FACS lysing solution according to the manufacturer’s instructions and leukocytes were analyzed by flow cytometry on a FACSCalibur using the BD Simulset software for data acquisition and analysis (Becton Dickinson). LDCs of validation set II and IV were determined either i) with an automated hematology analyzer (Coulter AcT diff2, Beckman Coulter, Brea, California, USA); ii) by microscopic analysis of blood smears; and/or iii) by immunophenotyping and flow cytometric analysis on a Navios flow cytometer (Beckman Coulter) to get proportions of T cells, CD4+ T cells, CD8+ T cells, B cells, and NK cells, as indicated in the text. Blood samples of validation set III were analyzed with an Abbott Cell-Dyn Emerald hematology system (Abbott Laboratories, North Chicago, Il, USA).

### Isolation of DNA and bisulfite conversion

Genomic DNA was isolated from blood with the QIAamp DNA Mini Kit (Qiagen, Hilden, Germany). Genomic DNA of serum was isolated from serum tubes (S-Monovette; Sarstedt, Nümbrecht, Germany) after one-hour incubation at room temperature and centrifugation for ten minutes at 2,000 x g. DNA was subsequently isolated from 1 ml serum with the PME free-circulating DNA extraction kit (GS/VL system; Analytik Jena, Jena, Germany) with addition of carrier RNA according to the manufacturer’s instructions. Either 1 μg of DNA from peripheral blood or the complete DNA sample from serum was bisulfite-converted with the EZ DNA Methylation Kit (Zymo Research, Irvine, CA, USA).

### Generation of non-methylated reference DNA for quantification

The target regions were PCR amplified (Eppendorf Mastercycler 5341; Eppendorf AG, Hamburg, Germany), cloned into the pBR322 vector (Thermo Fischer Waltham, Massachusetts, USA), expanded in DH5α *E.coli*, and isolated with the Plasmid DNA purification kit (Macherey-Nagel, Düren, Germany). To determine the optimal DNAm range of the blood-reference mixture, peripheral blood (150 μl) was mixed with serial dilutions of the reference sequence (0.0002 – 0.1100 ng). Mixtures of blood and reference DNA were subjected to DNA isolation and bisulfite conversion, as described above.

### Pyrosequencing

Specific regions covering the CpG sites cg05398700 (*WDR20*), cg17587997 (*FYN*), cg16452866 (*BCL11B*), cg05044173 (*CD4*), cg25939861 (*CD8A*), cg02665297 (*WIPI2*), cg13617280 (*SLC15A4*), cg10480329 (*CENPA*), cg06096175 (*LSM14B*), cg25834632 (*ZC3H3*), and cg09414987 (no associated gene) were amplified by PCR (Eppendorf Mastercycler 5341; Eppendorf AG). Primers were designed with the Pyrosequencing Assay Design Software 1.0 (Biotag AB, Uppsala, Sweden) and they are provided in Table S3. Pyrosequencing was then performed on a PyroMark Q96 ID System using region specific sequencing primers and results were analyzed with the PyroMark Q CpG software (Qiagen).

### MassARRAY analysis

Alternatively, we used MassARRAY for site-specific analysis of DNAm levels. Amplicons were designed with the Sequenom’s EpiDESIGNER software (Table S4). Converted DNA was amplified by PCR using the HotStart Plus PCR Master Mix (Qiagen). Unincorporated dNTPs were neutralized using shrimp alkaline phosphatase (Agena Bioscience, San Diego, CA, USA). Subsequently, 10 μl of PCR product was *in vitro* transcribed and cleaved in a base-specific (U-specific) manner using RNase A (T-Cleavage MassCleave Kit; Agena Bioscience, Hamburg, Germany). The cleaved products were then analyzed by the MALDI-TOF mass spectrometer (MassARRAY Analyzer 4 System; Agena Bioscience). Measurements were performed at Varionostic GmbH (<u>www.varionostic.de</u>; Ulm, Germany).

### Combination of DNAm levels for Epi-Blood-Count

Linear regression of DNAm levels at individual CpG sites was initially determined in 60 samples of the HELPcB training set. The regression formulas (indicated in Fig. S3) were then used to estimate cell counts in a validation set. Alternatively, we used a deconvolution approach that combines DNAm data of various cell types (three, five, or six CpGs). The Epi-Blood-Count can be represented by a matrix *W* of size f x k (f: number of CpGs (features); k: number of cell types). The methylation data of the blood samples are represented by a matrix *V* of size f x n (n: number of blood samples) and are modeled as a linear combination of the purified cell types *W*, with their mixture proportions *H* (k x n matrix – each of the n columns corresponds to the mixture proportion of the respective blood sample (same column in *V*)): *V ≈ WH*

For estimation of *H*, a non-negative least-squares (NNLS) approach is used to avoid negative mixture proportions. For implementation purposes, we use the multiplicative update rule of Lee et al. (Lee & Seung, 2001) to determine *H*:

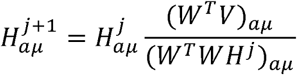

Here, *j* is the iteration index, *W^T^* indicates the transpose of matrix *W*, and *α* and *μ* are the row and column indices, respectively. For a unique estimation of *H*, it is required to have at least as many CpG sites as cell types, i.e. *f* ≥ *k* (assuming that the rows of *W* are linearly independent). Once leukocyte proportions were calculated, we added the proportions together and adjusted them to a total sum of 100%.

If no measurements from purified cell types are available, it is possible to use the reverse approach, estimating matrix *W* from *H* and *V*, given a set of peripheral blood samples with available cell counts (*H*) and methylation data (*V*). In order to do so, we use the respective iterative formula (Lee & Seung, 2001) for estimating *W*:

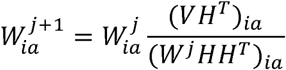

### Quantification of cell numbers based on DNAm levels

Upon mixture of genomic DNA with a non-methylated reference DNA the DNAm level can be mathematically described as ratio of methylated to total DNA:

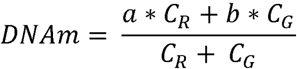

Here, *C_R_* and *C_G_* resemble the copy number of the reference DNA and the genomic DNA, respectively; *a* and *b* are absolute DNAm levels that were determined by pyrosequencing or MassARRAY in controls consisting of either pure reference DNA or blood DNA, respectively (e.g. 7% and 93% DNAm in our analysis). To determine the copy number of the reference plasmids (*C_R_*) we used the following formula:

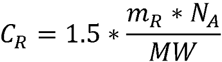

where *m_R_* is the added reference amount (e.g. 0.011 ng of *LSM14B*), *N_A_* is the Avogadro’s constant, MW is the molar weight of the reference DNA (calculated for the plasmid with *LSM14B* sequence: 2.85*10^6^ gmol^-1^), and the correction factor 1.5 was empirically determined for our preparation of reference DNA and relates to the fact, that purified plasmids may also comprise fragments of the bacterial genome or other plasmids.

With these parameters it is inversely possible to calculate copy numbers of the genomic DNA (*C_G_*) – and hence the cell numbers in a given blood sample:

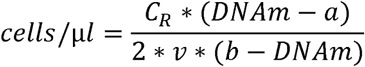

The term “2” stems from the fact that each cell comprises two copies of genomic DNA; *v* is the volume of analyzed blood in μl.

### Statistical Analysis

Mean average deviation (MAD), mean standard error (MSE), Pearson correlation coefficient (R), linear regression, and Student’s t test were calculated in Excel. *P* values < 0.05 were considered as statistically significant.

## Acknowledgements

This work was supported by the Else Kröner-Fresenius-Stiftung (2014_A193), by the German Research Foundation (WA 1706/8-1), and by the German Ministry of Education and Research (01KU1402B). M.L. was supported by the Ministry for Innovation, Science and Research of German Federal State of North Rhine-Westphalia, Germany; and the Dutch Province of Limburg, The Netherlands.

## Author contributions

JF and TB designed the study, performed pyrosequencing experiments, and analyzed the data. ML developed the NNLS algorithm and helped with computational analyses. CB and UG performed MassARRAY experiments. YH assisted with pyrosequencing experiments. PU, KS, SI, JP, AE, TK, LR and SK provided and characterized blood samples. RH performed manual blood counts. WW conceived the study and participated in data analysis, and coordination. WW, JF and TB wrote the manuscript and all authors read, edited, and approved the final manuscript.

